# Actin dynamics drive microvillar motility and clustering during brush border assembly

**DOI:** 10.1101/432294

**Authors:** Leslie M Meenderink, Matthew J. Tyska

## Abstract

During differentiation, transporting epithelial cells generate large arrays of microvilli known as a brush borders to enhance functional capacity. To develop our understanding of brush border formation, we used live cell imaging to visualize apical surface remodeling during early stages of this process. Strikingly, we found that individual microvilli exhibit persistent active motility, translocating across the cell surface at ~0.2 μm/min. Perturbation studies with inhibitors and photokinetic experiments revealed that microvillar motility is driven by actin assembly at the barbed-ends of core bundles, which in turn is linked to robust treadmilling of these structures. Because the apical surface of differentiating epithelial cells is crowded with nascent microvilli, persistent motility promotes collisions between protrusions and ultimately leads to their clustering and consolidation into higher order arrays. Thus, microvillar motility represents a previously unrecognized driving force for apical surface remodeling and maturation during epithelial differentiation.

## INTRODUCTION

Microvilli are evolutionarily ancient cell surface protrusions (Sebe-Pedros et al., 2013) that consist of a core bundle of parallel actin filaments enveloped in plasma membrane. Epithelial cells have adapted this simple structure to fulfill physiologically diverse functions including mechanosensation in the inner ear sensory stereocilia (Schwander et al., 2010), chemosensation in the lung, gut, and urogenital tract (Gerbe and Jay, 2016; Krasteva and Kummer, 2012; Kummer and Deckmann, 2017), and solute uptake in the renal tubules (Coudrier et al., 1988) and intestine (Delacour et al., 2016). Most epithelial cells assemble densely packed apical arrays of microvilli referred to as ‘brush borders’ based on their appearance. Disruption of brush border function through inherited defects (Schneeberger et al., 2018), infectious causes (Vallance et al., 2002), or injury (Emlet et al., 2015) leads to a range of human diseases.

Detailed studies on intestinal epithelial brush borders have revealed that microvillar cores contain 20-30 parallel actin filaments with barbed-ends, the preferred site of monomer addition, oriented toward the distal tips and pointed-ends anchored in a terminal web (Mooseker and Tilney, 1975). Actin cores are stabilized by bundling proteins including villin (Bretscher and Weber, 1979), espin (Bartles et al., 1998), fimbrin (also known as T-plastin) (Bretscher and Weber, 1980), and potentially Eps8 (Hertzog et al., 2010). Additionally, core bundles are linked to the overlying plasma membrane by myosins (Tyska et al., 2005) and ERM proteins (Casaletto et al., 2011). More recent studies indicate that microvillar growth and elongation are regulated by the WH2 domain protein Cobl and BAR domain protein IRTKS (Grega-Larson et al., 2015; Postema et al., 2018; Wayt and Bretscher, 2014).

Recent work also offers some insight on factors that organize microvilli in mature brush borders. We now know that microvillar packing is driven by an intermicrovillar adhesion complex (IMAC) that forms between the tips of adjacent protrusions. The IMAC is composed of the interacting cadherins CDHR2 (protocadherin-24) and CDHR5 (mucin-like protocadherin), which are positioned at the distal tips through an interaction with the cytoplasmic tripartite assembly of USH1C (Harmonin), ANKS4B, and MYO7B (Crawley et al., 2014b; Crawley et al., 2016; Weck et al., 2016). Ultrastructural analysis of Caco-2_BBE_ cells through stages of differentiation also revealed that individual microvilli first form small tepee-shaped clusters, which progressively increase in size (i.e. number of microvilli) until the array fills the entire apical surface, becoming a mature brush border (Crawley et al., 2014b). How adherent clusters of microvilli form and whether adhesion complexes form before or after microvillar growth remains unknown.

Despite growing knowledge and identification of factors that contribute to the assembly of microvilli, how cells organize these structures into mature brush borders remains poorly understood, primarily due to a lack of time-resolved data on apical surface remodeling during this process. Using spinning disk confocal microscopy (SDCM) to characterize the early steps of brush border maturation, we found that individual microvilli exhibit persistent, active motility. Microvillar movement is driven by actin assembly at microvillar tips, which in turn promotes robust treadmilling that propels protrusions across the cell surface. Although myosin contractility does not drive microvillar motility, it does control the length of motile structures. Importantly, we found that microvillar motility promotes collisions and clustering of individual protrusions as well as their consolidation into higher order arrays. This work identifies microvillar motility as a driving force for apical remodeling during epithelial differentiation.

## RESULTS

### Microvilli exhibit persistent, active motility

To examine the behavior of microvilli early in differentiation, we imaged the apical surface of LLC-PK_1_ clone 4 (CL4) cells, which are derived from porcine renal proximal tubules (Hull et al., 1976) and have been used in past studies of microvillar biology (Bartles et al., 1998; Tyska and Mooseker, 2002). Structured illumination microscopy (SIM) of CL4 cells one day post confluence (1 DPC) revealed large numbers of plasma membrane-wrapped linear actin bundles on the apical surface, consistent with nascent microvilli (Figure S1A-C). To gain insight on the dynamics of microvillar growth and brush border maturation, we used live cell SDCM to image cells expressing fluorescent protein-tagged espin, which targets specifically to the parallel actin bundles that support microvilli (Figure 1A). Nascent microvilli exhibit a continuum of orientations, ranging from parallel to perpendicular to the cell surface (Figure 1B). SDCM of CL4 cells co-expressing mCherry-Espin and GPI-GFP confirmed that microvilli in both orientations are enveloped in plasma membrane (Figure 1C, D), and that membrane wrapping persists over time (Supplemental movie 1). Of note, espin was selected for these studies because it localizes along the length of microvilli and has limited impact on actin turnover dynamics (Loomis et al., 2003). However, we did observe microvillar lengthening similar to that reported by others in cells expressing high levels of tagged espin. Therefore, for time-lapse imaging we selected cells expressing low levels of espin, and under these conditions maximum length was no different compared to cells expressing the F-actin probe, EGFP-Lifeact (Figure S1D and Figure 1J).

**Figure 1.**
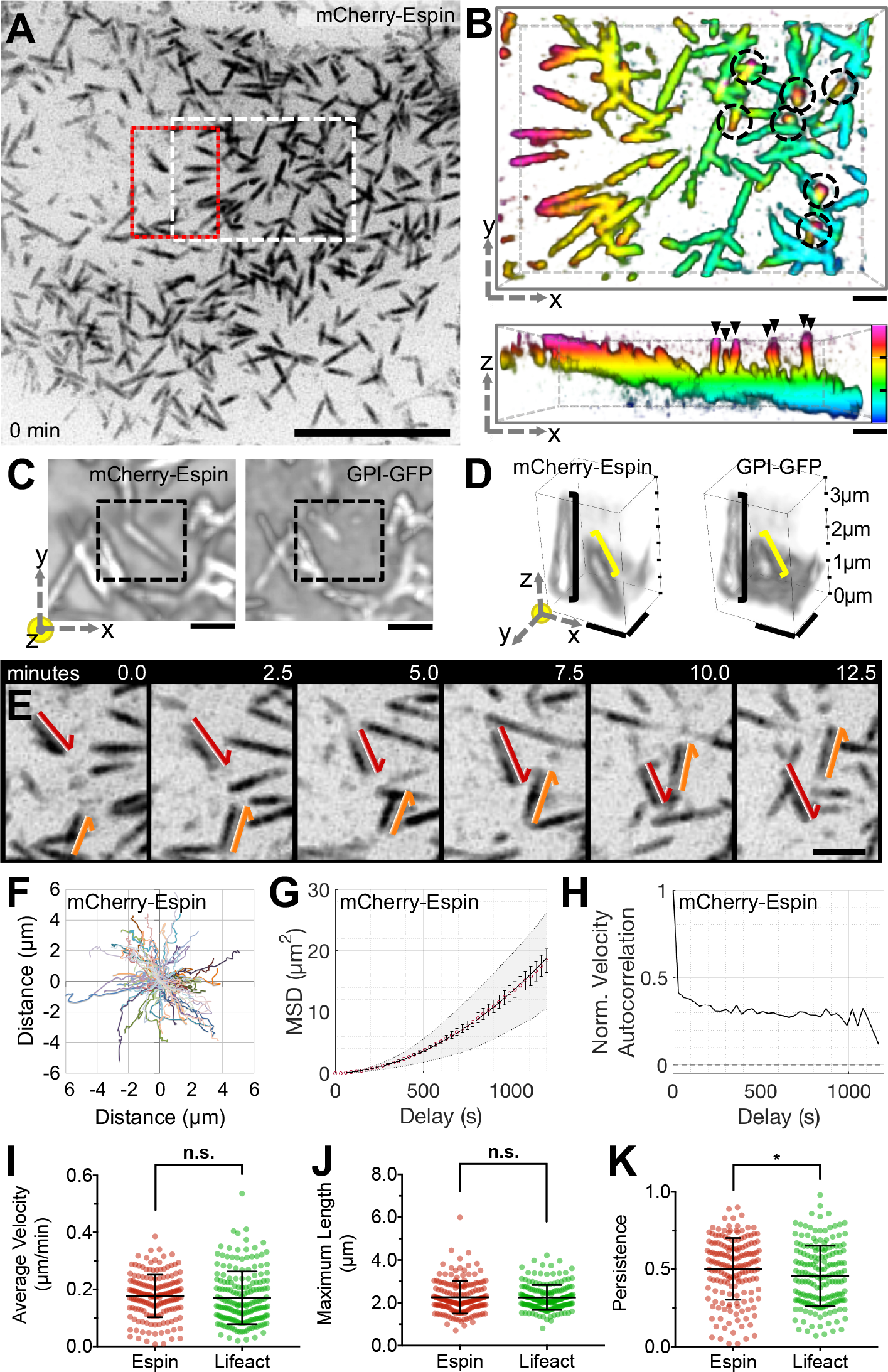
Microvilli exhibit persistent, active motility. (A) SDCM of the apical surface of a CL4 cell stably expressing mCherry-Espin, viewed as a maximum intensity z-projection. Scale bar 10 μm, white dashed box corresponds to B, red dashed box corresponds to E. (B) 3-dimensional (3D) depth color coded z-stack viewed *en face* (xy plane, upper panel) or laterally (xz plane, lower panel). Scale bars are 1 μm, z-axis depth color code (lower panel) to scale with tick marks at 1 μm intervals. Microvilli exhibit a range of orientations from parallel (represented by a single color) to perpendicular to the cell surface (spanning multiple color bands, circled—top panel, arrowheads—bottom panel). (C) SDCM of a CL4 cell expressing mCherry-Espin and GPI-GFP. A single z-stack was deconvolved and viewed as a 3D alpha-blended composite. Scale bar is 1 μm. (D) Volume from boxed area in C was cropped and rotated to emphasize two membrane-wrapped microvilli. One microvillus extends ~3 μm from the cell surface (black bracket), the other shorter structure is nearly parallel to the cell surface (yellow bracket). Brackets indicate the extent of the membrane signal. Scale bars are 1 μm. (E) Time series of microvilli translocating across the cell surface; red and orange arrows highlight the paths of two distinct protrusions. Scale bar 2 μm. (F) Rose plot of trajectories measured from the tips of microvilli (n = 101) for the cell in A. (G) Microvillar trajectories from F were subject to MSD analysis; red open circles represent the mean MSD, error bars indicate standard error of the mean (SEM), grey area marks the weighted standard deviation (SD) over all MSD curves, and the solid line indicates a fit of the data to an active movement model (diffusion coefficient, D = 0.000283 μm^2^/s and velocity, V = 0.21 μm/min). (H) Trajectories from F were analyzed for normalized velocity autocorrelation, solid line. Dotted line at 0 indicates the velocity autocorrelation of random diffusive movement. (I-K) Average microvillar velocity, maximum microvillar length, and persistence, respectively. Error bars indicate mean ± SD, n = 171 microvilli from 7 cells, * p < 0.05, n.s. = not significant.

Strikingly, SDCM of CL4 cells revealed that microvilli are highly motile with actin core bundles translocating across the cell surface (Figure 1E, Supplemental Movie 2) at a mean velocity of 0.18 μm/min (Figure 1I). This rate is similar to microvillar growth rates reported for *Xenopus* kidney epithelial A6 cells and mouse cells cultured from the inner ear (Gorelik and Gautreau, 2014), but is substantially lower than rates reported for filopodial extension (≥1 μm/min) (Mallavarapu and Mitchison, 1999). To investigate potential patterns of motility, we tracked large numbers of protrusions. Rose plot analysis of the resulting trajectories did not reveal obvious flow patterns or points of nucleation (Figure 1F). However, mean square displacement (MSD) analysis generated a parabolic curve consistent with active motility (Figure 1G). This was further supported by a positive velocity autocorrelation function (Figure 1H) where purely diffusive movement has a velocity autocorrelation of 0 and active movement greater than 0. Notably, microvilli exhibited similar motile properties in cells expressing EGFP-Lifeact (Figure S1D-H and Figure 1I-K), although our remaining experiments took advantage of the specificity and enhanced signal/noise offered by the espin probe.

### Microvillar motility is driven by actin assembly

Motility of subcellular structures can be driven by diffusion, cytoskeletal polymerization, or molecular motors. Because microvillar motion is not diffusive (Figure 1G, H), we focused on testing the involvement of actin assembly and myosin-2 powered contractility. Based on brush border ultrastructure (Mooseker and Tilney, 1975), both mechanisms are relevant and could contribute to microvillar movement. To dissect the contribution of these activities, we used inhibitors to block myosin-2 contractility (Blebbistatin) or barbed-end actin assembly (Cytochalasin). CL4 cells were imaged as described above and inhibitors were added after 5 minutes (Supplemental Movie 3). After addition of 20 μM Blebbistatin, microvilli continued to translocate across the cell surface (Figure 2A-E). Trajectory analysis of microvilli on blebbistatin-treated cells showed a lower velocity (Figure 2K), but the overall pattern of motion, MSD plots, and autocorrelation outputs were consistent with controls (Figure 2B-E). Interestingly, maximum microvillar length increased significantly during Blebbistatin treatment (Figure 2L), which suggests that under normal conditions, myosin-2 may serve to limit the length of these protrusions. In contrast, microvillar translocation halted immediately after addition of Cytochalasin (Figure 2F-J). For detailed trajectory analysis, we used 500 nM Cytochalasin B, which had less impact on overall cell shape and facilitated tracking of individual microvilli. Trajectory analysis showed that these microvilli no longer exhibited persistent motion (Figure 2M), although they continued to exhibit slow, short range displacements over small areas (Figure 2G,H,K). MSD and autocorrelation analysis indicated behavior consistent with diffusion constrained around a point (Figure 2I,J). Together these data indicate that actin assembly at the barbed-ends of microvillar core bundles is required for the active motility of these structures across the apical surface.

**Figure 2.**
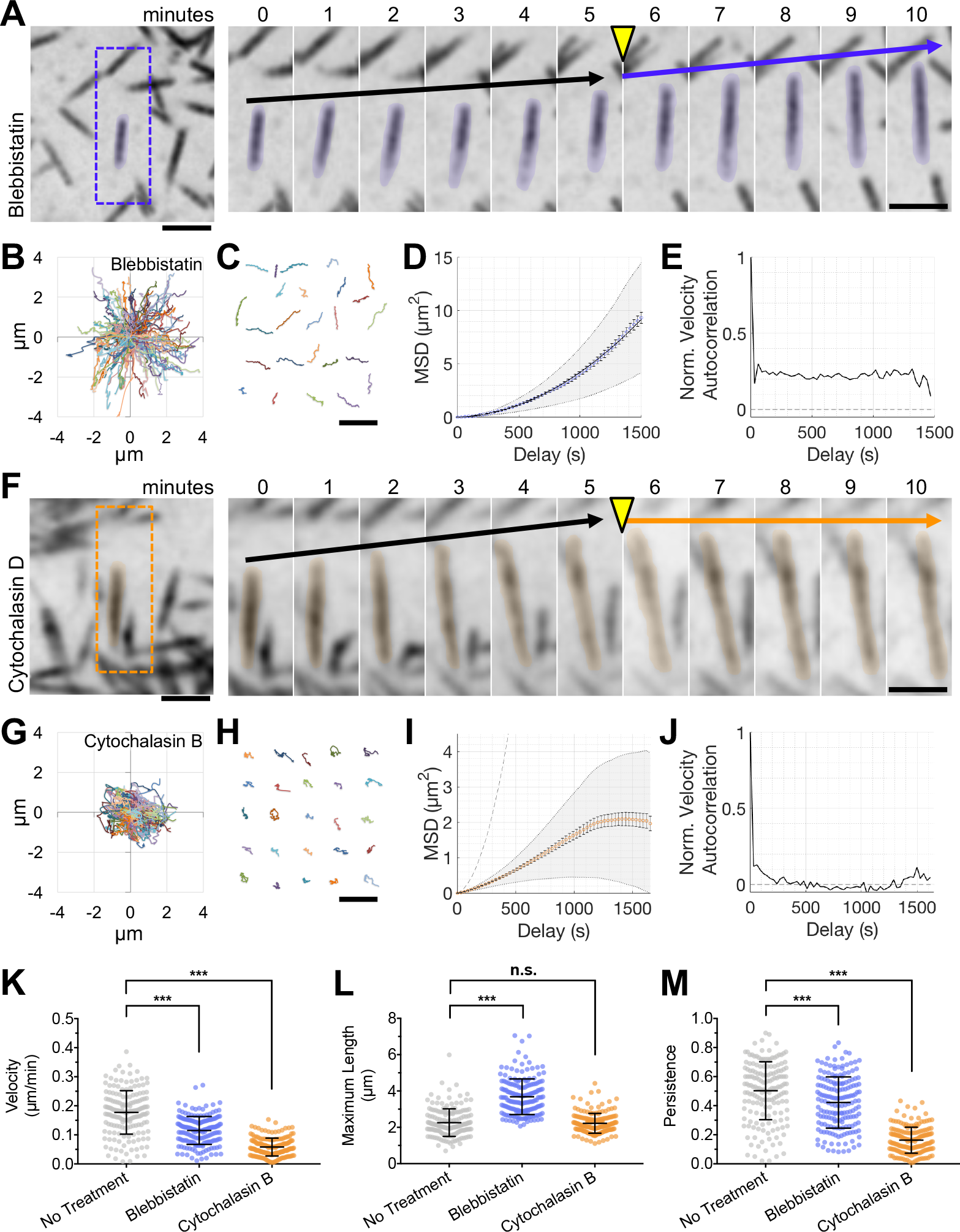
Microvillar motility is driven by actin assembly and is not dependent on myosin contractility. (A) SDCM of the apical surface of CL4 cells stably expressing mCherry-Espin showing the response to 20 μM Blebbistatin. *Right*, time series montage of a single protrusion highlighted with a 10% pseudo-colored overlay. Scale bars are 2 μm. Drug was added following the 5-minute time interval, yellow arrowhead. Black arrows indicate the baseline rate of microvillar movement. Blue arrow indicates the rate of microvillar movement after the addition of drug. (B) Rose plot shows the microvillar trajectories (n = 100) from a single cell treated with 20 μM Blebbistatin. (C) 25 representative microvillar trajectories are isolated for display. Scale bar is 5 μm. (D) MSD analysis of microvillar trajectories from B. (E) Normalized velocity autocorrelation analysis of microvillar trajectories from B; data were fit to an active movement model with D = 0.000093 μm^2^/s, V = 0.12 μm/min. (E) Normalized velocity autocorrelation analysis of microvillar trajectories from B. (F) SDCM of the apical surface of CL4 cells stably expressing mCherry-Espin showing the response to 30 μM Cytochalasin D. *Right*, time series montage of a single protrusion highlighted with a 10% pseudo-colored overlay. Scale bars are 2 μm. Drug was added following the 5-minute time interval, yellow arrowhead. Black arrows indicate the baseline rate of microvillar movement. Orange arrow indicates the rate of microvillar movement after the addition of drug. (G) Rose plot shows the microvillar trajectories (n = 100) from a single cell treated with 500 nM Cytochalasin B. (H) 25 representative microvillar trajectories are isolated for display. Scale bar is 5 μm. (I) MSD analysis of microvillar trajectories from G; data could not be fit with an active movement model. The curve for mCherry-Espin with no drug treatment is plotted for comparison (grey dotted line). (J) Normalized velocity autocorrelation analysis of microvillar trajectories from G. (K-M) Average microvillar velocity, maximum microvillar length, and persistence, respectively, measured from untreated cells (from Figure 1), or cells exposed to Blebbistatin or Cytochalasin B. For Blebbistatin and Cytochalasin datasets, n = 178 and 175 microvilli, respectively, from 6-7 cells. Bars represent mean ± SD. *** p < 0.0001, n.s. = not significant.

### Microvillar F-actin cores undergo treadmilling during motility

The robust inhibition of microvillar movement by Cytochalasin suggests a model where actin incorporation at the distal tips may provide a driving force for the translocation of microvilli. However, because microvilli maintain a relatively stable steady-state length, any barbed-end actin incorporation must be matched by disassembly from the pointed-ends of core bundles. Such a “treadmilling” process has been observed in microvilli before (Loomis et al., 2003; Tyska and Mooseker, 2002), although its relationship to microvillar motility has not been explored. To determine if microvillar core bundles exhibit treadmilling during motility, we expressed mNEON-Green-β-actin in CL4 cells to allow for direct monitoring of actin turnover. Cells were imaged using SDCM as before (Figure 3A), however, in this experiment we also photobleached fiduciary marks within translocating microvilli. To visualize the fate of fiduciary marks, images were then processed using Imaris to create three-dimensional (3D) surfaces representing apical microvilli (Figure 3B-D, see Supplemental Movie 4). We found that after creating a fiduciary bleach mark at time 0, microvilli continue to translocate over the cell surface. However, we noted a marked expansion of the actin signal at leading end of the protrusion (distal to the bleach mark) up to 1.1 μm/min, which was matched by a gradual loss of signal at the trailing end (Figure 3E). This pattern of turnover indicates that microvillar actin cores treadmill as they translocate across the cell surface with their barbed-ends in the lead.

**Figure 3.**
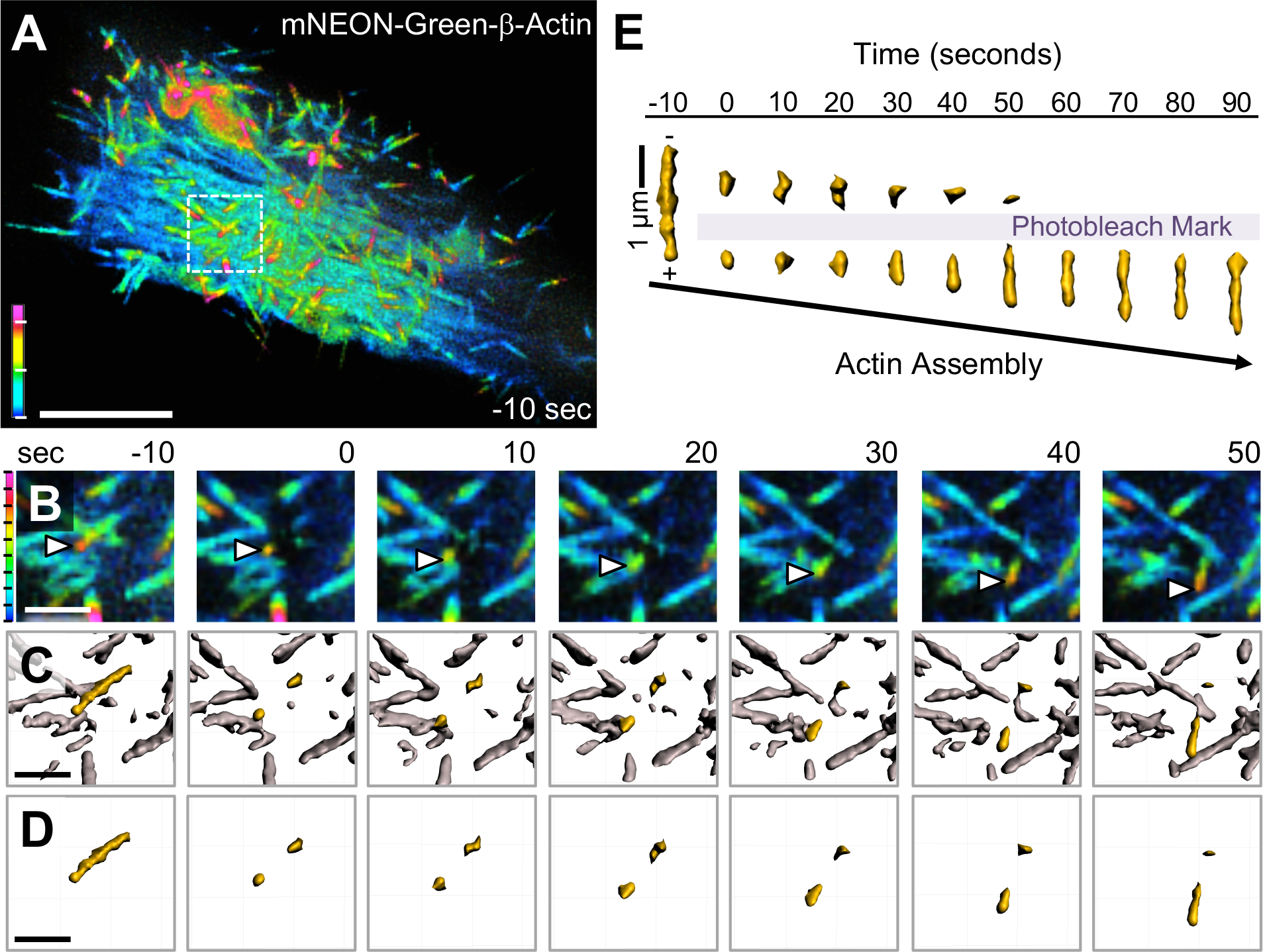
Microvillar F-actin cores undergo treadmilling during motility. (A) SDCM of a CL4 cell expressing mNEON-Green-β-actin viewed as a depth-coded z-projection. Scale bar is 10 μm, z-axis depth code with tick marks at 1 μm intervals is shown at lower left. (B) Time series montage of area in A highlighted by white dashed box, enlarged, cropped in z to remove cytoplasmic signal, then viewed as a depth coded z-projection. Time series shows region of interest before photobleaching (−10 seconds), immediately after photobleaching (0 seconds), and during recovery. White arrowheads mark the growing distal tip of a microvillus. (C) Data from B were processed using Imaris to create a 3D surface of the fluorescent actin signal with the microvillus of interest highlighted in yellow. (D) Time series montage showing the isolated microvillus of interest before and after photobleaching. In B-D, scale bar is 2 μm, z-axis depth code (left) with tick marks at 200 nm intervals. (E) The microvillus of interest from D was viewed orthogonally over time and aligned based on the position of the bleached region to highlight treadmilling of the mark through the actin core. Shown here is a representative example of treadmilling observed from n > 20 cells.

### Microvillar motility promotes intermicrovillar collisions, adhesion, and cluster formation

To understand if microvillar translocation contributes to organizing protrusions into a mature brush border, we studied mCherry-Espin-expressing CL4 cells later in differentiation (2 DPC). Specifically, we selected cells with a mixed population of individual microvilli and small clusters of protrusions. Imaging cells over several hours revealed that microvilli continue to translocate and are highly dynamic even at this later time point (Figure 4A, Supplemental Movie 5). These time-lapse data also revealed that microvillar motility promotes collisions between neighboring protrusions and these encounters can result in the formation of small clusters (Figure 4B, Supplemental Movie 6). Remarkably, even groups of microvilli in adherent clusters retain their ability to translocate across the cell surface, eventually colliding with other clusters and consolidating their numbers (Figure 4C, Supplemental Movie 7). These live cell observations indicate that microvillar motility provides the driving force for intermicrovillar adhesion and microvillar clustering during brush border maturation.

**Figure 4.**
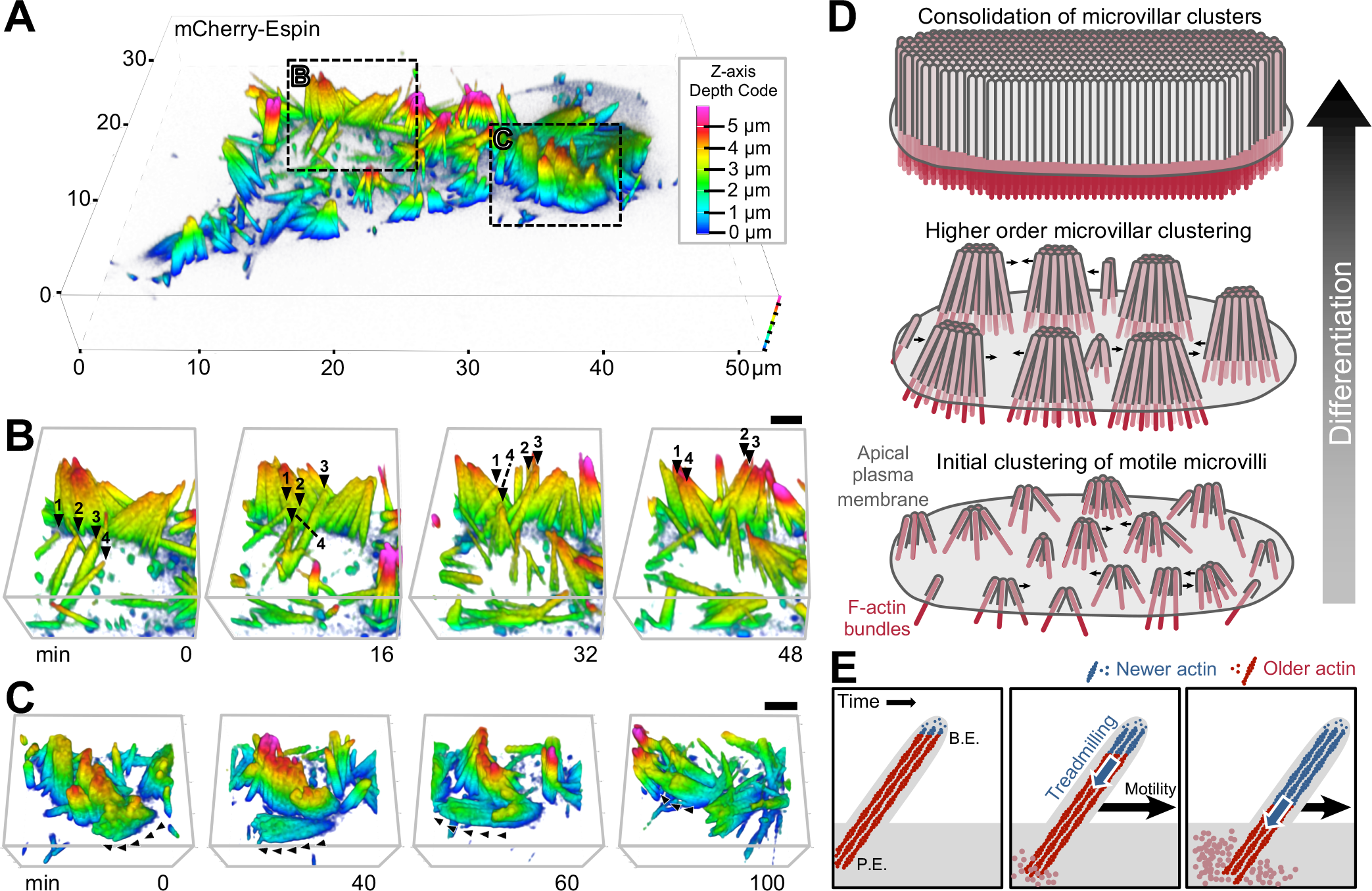
Microvillar motility promotes intermicrovillar collisions, adhesion, and cluster formation. (A) SDCM of the apical surface of a CL4 cell stably expressing mCherry-Espin at 2 DPC, visualized as a depth-coded composite. Image scale is shown on volume frame, depth scale is shown to the right. (B,C) Times series montages of areas highlighted in A, enlarged and rotated to optimize visualization of microvillar movement. In B, individual microvilli of interest are marked at each time point with numbered black arrowheads. In C, black arrowheads mark the path of motion for a cluster of microvilli. Scale bars are 2 μm. Z-axis depth code shown in A also applies to B and C. (D) A progressive clustering model for microvillar remodeling and brush border formation during differentiation. Here, nascent microvilli emerge from the apical surface and undergo persistent motility, which promotes collisions between protrusions. These encounters allow adhesion links to form between microvillar tips, resulting in characteristic tepee-shaped structures (originally reported in Crawley et al. *Cell* 2014). As maturation proceeds, clusters grow by moving across the apical surface, colliding with other clusters and consolidating their numbers until eventually the entire surface is occupied by one continuous large-scale cluster, i.e. a mature brush border. (E) Enlarged view of a single motile microvillus. The microvillar core is comprised of bundled F-actin (red) with the barbed-ends oriented toward the distal tip. New actin monomers incorporate at the barbed-ends (B.E., blue F-actin), which is balanced by monomer disassembly from the pointed-ends (P.E.). This results in ‘treadmilling’ of actin through the microvillar core which provides a pushing force against the membrane that powers microvillar motility.

## DISCUSSION

Here we report that microvilli undergo active movement across the apical surface of differentiating epithelial cells. Microvillar motion is driven by actin assembly at the barbed-ends of core bundles and linked to treadmilling of the bundle as structures translocate. Although previously unreported, this robust activity is harnessed by differentiating epithelial cells to organize and consolidate nascent microvilli into tightly packed arrays. Indeed, our time-lapse data provide clear evidence that active motility forces collisions between protrusions, resulting in the formation of larger microvillar clusters. These data also indicate that nascent microvilli do not emerge from the apical surface in adherent assemblies, but rather elongate from the surface and then interact following their initial growth to form adherent clusters. By identifying a mechanism that serves to initiate and amplify the formation of microvillar clusters, these discoveries provide strong support for the progressive clustering model proposed previously (Crawley et al., 2014a) and refine our understanding of the activities that culminate in a mature brush border (Figure 4D).

Actin filaments exhibit distinct assembly/disassembly kinetics at their two ends, with the barbed-ends being the preferred site for assembly (Pollard and Cooper, 2009). When monomer incorporation at the barbed-end is balanced by disassembly at the pointed-end, subunits can treadmill through the filament while a steady length is maintained. Polymer treadmilling is a well-established mechanism for generating mechanical force in diverse cellular processes ranging from chromosome separation in mitosis (Mitchison, 1989), to leading edge protrusion in cell motility (Wang, 1985), and pathogen movement during infection (Pantaloni et al., 2001). Actin monomer insertion at the barbed-end of a single filament can produce force in the piconewton range (Footer et al., 2007; Prass et al., 2006), deforming associated membrane structures (Giardini et al., 2003; Upadhyaya et al., 2003). With their exposed barbed-ends enveloped in plasma membrane, microvillar actin bundles are ideally positioned to generate a protrusive force at microvillar tips (Figure 4E). Our data show clear treadmilling of actin cores in translocating microvilli, linking this mechanism of force generation to microvillar motility.

While our studies indicate that myosin-2 is not essential for powering the motility of microvilli, the elongation of these structures after Blebbistatin application suggests that this motor plays a role in regulating their steady-state length. Myosin-2 has been localized to the terminal web (Bretscher and Weber, 1978) and previous studies implicate this motor in regulating microvillar orientation (Temm-Grove et al., 1992). The mechanism driving microvillar elongation under these conditions remains unclear, but myosin may assist with turnover of F-actin at the pointed-ends, which would be reminiscent of contractility-driven actin turnover observed in other systems (Medeiros et al., 2006; Wilson et al., 2010). Future studies examining the organization of terminal web myosin-2 and its attachment to microvillar actin cores will be needed to fully appreciate the role of this motor in microvillar dynamics and brush border assembly.

## ACKNOWLEDGEMENTS

We thank all members of the Tyska laboratory for feedback and advice. Microscopy was performed in part through the Vanderbilt University Cell Imaging Shared Resource. This work was supported by a Ruth L. Kirschstein National Research Service Award T32-A1007474 (LMM) and NIH grants R01DK111949 and R01DK095811 (MJT).

## AUTHOR CONTRIBUTIONS

MJT conceived the study. LMM and MJT designed experiments. LMM performed all experiments, analyzed all data, and prepared the figures. LMM and MJT wrote the manuscript.

## METHODS

### Cell Culture and Stable Cell Line Generation

LLC-PK_1_-CL4 (CL4) cells were cultured at 37°C and 5% CO_2_ in DMEM with high glucose and 2 mM L-glutamine supplemented with 10% fetal bovine serum (FBS). For generation of stable cell lines CL4 cells were grown to 80-90% confluency in T25 flasks and transfections were performed using Lipofectamine 2000 (Invitrogen) or FuGENE 6 (Promega) according to the manufacturer’s instructions. Selection for stable expression was performed after cells recovered for 2-3 days with addition of 1 mg/ml G418.

### Constructs

The pmCherry-Espin construct was a kind gift from Dr. James Bartles. The pGL-GPI-GFP was provided by the Dr. Anne Kenworthy (University of Virginia). The EGFP-Lifeact construct was provided by Dr. Irina Kaverina (Vanderbilt University). The mNEON-green-β-actin was purchased from Allele Biotechnology.

### Microscopy and Image Processing

For SIM imaging, cells were plated on glass coverslips and allowed to grow to confluence. Cells were washed with warmed phosphate-buffered saline (PBS) and fixed with warm 4% paraformaldehyde/PBS for 15 min at 37°C. Cells were then washed three times with PBS, and blocked for 1 hour at room temperature in 5% bovine serum albumin (BSA) in PBS. Alexa Fluor 568-phalloidin (1:200, A12380; Invitrogen) or WGA-488 (10 μg/ml; W11261 ThermoFischer Scientific) were diluted in blocking solution and incubated for 1 hour at room temperature. Coverslips were washed three times with PBS then mounted on glass slides in ProLong Gold (P36930; Invitrogen). Cells were imaged on a Nikon Structured Illumination Microscope (N-SIM), Andor DU-897 EMCCD camera with four color excitation lasers (405 nm, 488 nm, 561 nm, and 647 nm), and a 100x/1.49 NA TIRF objective. SIM images were reconstructed using the Nikon Elements reconstruction algorithm. For live-cell SDCM, cells were plated onto plasma-cleaned 35 mm glass bottom dishes (Invitro Scientific, D35-20-1.5-N), then transfected with the appropriate marker construct. Cells were allowed to grow to the appropriate level of confluence. If transfected, cells were imaged within 24 to 72 hours of transfection. Live-cell imaging was performed on a Nikon Ti2 inverted light microscope equipped with a Yokogawa CSU-X1 spinning disk head, Andor DU-897 EMCCD camera or a Photometrics Prime 95B sCMOS camera, 488 nm and 561 nm excitation lasers, a 405 nm photo-stimulation laser directed by a Bruker mini-scanner to enable targeted photoactivation, photoconversion, and photobleaching), and a 100x/1.49 NA TIRF objective. Low density microvilli were imaged when cells were subconfluent (80-90% confluence) or 1 DPC and images were acquired every 30-60 seconds for 20-40 minutes (Figures 1 and 2, and Supplemental Figure 2). During drug treatment, either (-)-Blebbistatin (B592500 Toronto Research Chemicals), Cythochalasin D (C2618 Sigma), or Cytochalasin B (C6762 Sigma) was added after 5 minutes of baseline measurement (Figure 2). For photokinetic studies of actin dynamics, baseline images were obtained for several frames prior to bleaching, then for an additional 3-5 minutes of recovery at 10 second intervals. Bleaching was performed on a line ROI (0.1 μm in width and 3-15 μm in length) using a 405 nm laser at 30% power with a 10 μs dwell time. Higher density microvilli on cells 2 DPC were imaged every 1-2 minutes for up to 4 hours (Figure 4). During imaging, cells were maintained with humidity at 37°C with 5% CO2 using a stage-top incubation system. Image acquisition was controlled with Nikon Elements software. 3D time series images were oversampled in the z-dimension with z-steps ranging from 0.09 μm to 0.18 μm followed by deconvolution (Nikon Elements Automatic or Richardson-Lucy algorithms) for better object resolution. Images were contrast enhanced, cropped, and aligned using Image J software (NIH), Nikon Elements, or Imaris (BITPLANE). Two-dimensional images were viewed as a maximum intensity projection. Three-dimensional depth coding was completed using Nikon Elements with images viewed as alpha-blended 3D composite images. Imaris (BITPLANE) was used to create an initial surface representing the microvillar fluorescence signal (Figure 3) then manually adjusted with fusion/fission of adjacent objects and manual microvillar tracking.

### Quantification, Calculations, and Statistical Analysis

Microvillar centroids (x and y coordinates, length, and angle) were manually tracked using ImageJ, and the tip position was calculated using Microsoft Excel (Figure 1F, 2C-D, 2G-H, and Supplemental Figure 2C). The rate of microvillar motility was calculated as the net microvillar tip movement over microvillar lifetime. Maximum microvillar length is the longest length measured for a microvillus during its lifetime. Persistence is calculated as the net microvillar tip displacement divided by the total path traveled. If tips move in a straight line, persistence is equal to 1. All calculations on drug treated samples use the time of drug addition as time = 0. Mean square displacement (MSD) and velocity autocorrelation analysis were completed on microvillar centroids tracked in ImageJ. Data was exported and analyzed with MATLAB using a package specifically developed for MSD analysis (Tarantino et al., 2014), which is publicly available at http://www.mathworks.com/matlabcentral/fileexchange/40692-mean-square-displacement-analysis-of-particles-trajectories. For each individual trajectory, the MSD as a function of time delay (seconds) was calculated using 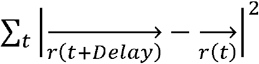, and then averaged overall all trajectories. Random motion is indicated by a linear relationship whereas active motility is represented by convex curvature, and restricted motion by concave curvature. Active motility was fit to *MSD*(*n*) = 4*Dn* + *v*^2^*n*^2^, where D is the diffusion coefficient, V is the velocity, and n is the time window used for calculation. For MSD plots, open circles represent the mean MSD, error bars are the SEM, and shaded areas represent the weighted SD over all MSD curves. The velocity autocorrelation is likewise calculated for each individual track and then averaged over trajectories. For random motion, displacements are not correlated, thus velocity autocorrelation is 0 except for the initial positive displacement (normalized to 1). For active motion, non-zero velocity autocorrelation is expected. For all figures, error bars indicate SD, and n values are reported in the figure legends. Statistical significance was calculated using the Mann-Whitney-Wilcoxon test for pairwise comparisons (Figure 1I-J), or using one-way ANOVA with Tukey’s multiple comparison to compare more than 2 conditions (Figure 2K-M). Rose plots and individual tracks were graphed using Excel. MSD and normalized velocity autocorrelation were analyzed and graphed using MATLAB. All other graphs were generated in Prism v.7 (GraphPad). All statistical testing was performed in Prism v.7 (GraphPad) or MATLAB.

## SUPPLEMENTAL INFORMATION

**Figure S1. Visualization and characterization of nascent CL4 cell microvilli.** (A) Fixed CL4 cell stained with Phalloidin (magenta) and wheat germ agglutinin (WGA, green) to mark F-actin and membrane, respectively. Boxed areas correspond to B. Scale bar is 10 μm. (B) Enlarged images highlight individual microvilli with distal tip membrane coverage (i) or more complete membrane coverage (ii). Scale bar is 1 μm. (C) Line scan intensity plots of phalloidin and WGA signal along the microvillar axis; arrows in B indicate line scan position and orientation. (D) SDCM of the apical surface of a CL4 cell transiently expressing EGFP-Lifeact. Scale bar is 10 μm. (E) Time series montage of boxed area in D; arrows highlight a microvillus translocating across the cell surface. Scale bar is 2 μm. (F) Rose plot of trajectories calculated from the tips of microvilli from D (n = 102). (G) Microvillar trajectories from F were subject to MSD analysis; data were well fit to an active movement model (black line) where D = 0.000361 μm^2^/s and V = 0.14 μm/min. (H) Microvillar trajectories from F were subject to normalized velocity autocorrelation analysis. The dotted line at zero indicates the velocity autocorrelation for random diffusive movement. Similar trajectories, MSD and autocorrelation results were observed across replicate cells (n = 81 microvilli from 7 cells).

**Supplemental Video 1. Early actin structures protruding from the cell surface are wrapped in membrane over time.** Time lapse images of CL4 cell stably expressing mCherry-Espin (left panel) and GPI-GFP (right panel), corresponds to Figure 1C-D. The 3D volume was rotated so the microvillus of interest is central and moves vertically across the image, and a depth-code was applied. Z-axis depth color code bottom left with tick marks at 1 μm intervals. Movie contains 6 frames taken at 1 minute intervals.

**Supplemental Video 2. Microvilli move across the cell surface.** Time lapse images of CL4 cell stably expressing mCherry-Espin (left panel, corresponds to Figure 1A), or EGFP-Lifeact (right panel, corresponds to Figure S1D). Duration 20 minutes, 12 frames per second (fps). Color-coded arrowheads highlight individual translocating microvilli. Scale bars are 10 μm.

**Supplemental Video 3. Microvillar motility requires actin assembly but not myosin contractility.** Time lapse images of CL4 cell stably expressing mCherry-Espin treated with 20 μM Blebbistatin (left panel, corresponds to Figure 2A) or 500 nM Cytochalasin B (right panel, corresponds to Figure 2F) with the addition of drug at 5 minutes. Duration 30 minutes, 12 fps. Scale bars are 10 μm.

**Supplemental Video 4. Microvillar F-actin core treadmills during microvillar motility.** Time lapse images of CL4 cell transiently expressing mNEON-Green-β-actin (top panel), microvillar surface created in Imaris (middle panel), isolated microvillus of interest (bottom panel). Images correspond to Figure 3B-D. Duration 120 seconds, 5 fps. Scale bars are 2 μm.

**Supplemental Video 5. Microvillar translocation drives intermicrovillar collision facilitating cluster formation.** Time lapse images of CL4 cell stably expressing mCherry-Espin. Duration 120 minutes, 10 fps. Movie corresponds to Figure 4A.

**Supplemental Video 6. Microvillar translocation drives collision and clustering of individual microvilli.** Time lapse images of CL4 cell stably expressing mCherry-Espin. Individual microvilli collide and remain connected then translocate across the cell joining with larger clusters of microvilli. Single microvilli are marked by arrowheads, then asterisks once they coalesce with a larger cluster. Duration 120 minutes, 10 fps. Movie corresponds to Figure 4B, zoom of Supplemental Movie 5.

**Supplemental Video 7. Large clusters of microvilli move across the cell surface.** Time lapse images of CL4 cell stably expressing mCherry-Espin. A large cluster of microvilli (asterisk) moves across the cell surface traveling left then upward. Duration 120 minutes, 10 fps. Movie corresponds to Figure 5C, zoom of Supplemental Movie 5.

